# Proteomic analysis of bronchoalveolar lavage fluid after lung transplantation associates stable allograft function with less lung damage at 12 months

**DOI:** 10.1101/2025.06.08.658472

**Authors:** Liisa Arike, Kristina Johansson, Anna Ermund, Mark Greer, Thaher Pelaseyed, Johan Westin, Gunnar C Hansson, Jesper M Magnusson

**Affiliations:** Department of Medical Biochemistry and Cell Biology, Institute of Biomedicine, University of Gothenburg, Gothenburg, Sweden; Department of Internal Medicine and Clinical Nutrition, Institute of Medicine, University of Gothenburg, Gothenburg, Sweden; Department of Respiratory Medicine & Infectious Diseases, Hannover Medical School, Hannover, Germany; German Centre for Lung Research (DZL/BREATH), Hannover, Germany; Department of Infectious Diseases, Institute of Clinical Sciences, Sahlgrenska University Hospital, Gothenburg, Sweden; Department of Respiratory Medicine, Institute of Medicine, Sahlgrenska University Hospital, Gothenburg, Sweden

## Abstract

**Introduction:** Freedom from chronic lung allograft dysfunction (CLAD) is a key objective after lung transplantation, yet predicting its onset remains challenging. This study investigated whether early proteomic changes in bronchoalveolar lavage fluid (BALF) can differentiate between patients maintaining stable graft function at 36 months and those developing CLAD within the first year. Additionally, findings were compared to proteomic data from non-transplanted individuals.

**Methods:** BALF samples were collected at one and twelve months post-transplant from 43 lung transplant recipients together with clinical parameters. Proteomic analysis was performed using mass spectrometry with label-free quantification for global protein profiling and heavy-labelled peptides for absolute quantification of mucins and related proteins. Differentially expressed proteins were identified and analyzed through pathway enrichment to explore biological mechanisms associated with CLAD.

**Results:** No significant proteomic differences were detected at one month. By twelve months, 63 proteins were differentially expressed between patients who developed early CLAD and those with stable function. Mucin levels declined in stable patients but remained elevated in both groups compared to healthy controls. Cartilage acidic protein 1 was significantly higher in stable patients at twelve months and correlated with better pulmonary function. Pathway analysis linked several altered proteins in CLAD patients to networks associated with lung injury and remodelling.

**Conclusion:** Protein profiles in BALF that resemble those of healthy lungs are associated with sustained graft function, while persistent expression of lung injury markers is associated with early CLAD. This suggests an adaptive process is needed for long-term post-transplant success.

## Introduction

Lung transplantation (LTx) is a key treatment for end-stage, nonmalignant pulmonary disease when other treatments have failed (1). Despite its history of over 60 years (2), the procedure still has worse survival than other solid organ transplants pioneered in the same era (3). Chronic lung allograft dysfunction (CLAD) is the major cause of death after the first post-operative year, affecting about 50% of recipients within five years (4). Current understanding of CLAD pathophysiology involves complex interactions of the innate and adaptive immune system and structural tissue remodeling due to repeated lung injury, ultimately leading to allograft failure (5). Thus, early post-operative CLAD is a major clinical setback, and given the post-operative clinical complexities, the underlying mechanisms are convoluted.

LTx presents unique challenges compared to other solid organ transplants due to environmental exposure to particles, pathogens, toxins, and allergens. These interact with the allograft in an immunosuppressed host, increasing the risk of respiratory infections associated with CLAD (6). Alongside other airway defense mechanisms, the mucus clearance system is essential in protecting the lungs against harmful microbes and infections (7). Under normal circumstances, ciliary beating transports bundles of mucus strands from submucosal glands and mucus threads from surface cells in an efficient system for moving foreign material out of the lung, keeping the lungs essentially clean (8–10). Following infection, mucin secretion, especially MUC5AC is promoted (11). An attached mucus layer is formed, like other chronic lung diseases such as chronic obstructive pulmonary disease (COPD) (12). The status of the mucociliary clearance after lung transplantation is likely abnormal, as indicated by reduced mucociliary clearance at least three months after LTx (13). Also, the transplanted lung is denervated from the lower vagal nerve, which postoperatively impairs the cough reflex for a period of six to twelve months (14). To what extent the transplanted lung can mimic normal function after transplantation and how the transplantation outcome is reflected in the lung proteome, particularly mucin-related proteins, is unknown.

In this observational exploratory study of LTx recipients, patients developing CLAD within the first-year post-transplant (early CLAD; eCLAD) are compared to those alive with stable lung function for at least three years. Using mass spectrometry (MS), protein abundance in bronchoalveolar lavage fluid (BALF) at one (1M) and twelve months (12M) after transplantation is compared. The proteomic results were also compared to healthy non-smokers, healthy smokers and COPD patients (12) using same heavy-labeled peptides. MS analysis was combined with BALF cellularity and pulmonary function tests (PFT) to identify potential mediators associated with outcomes after LTx.

## Material and methods

### Study design

Two patient groups were established from an existing cohort with intensified infection monitoring (15): LTx recipients exhibiting stable lung function in the first 36 months post-LTx (Stable) and patients who developed CLAD within a year post-LTx (eCLAD). Sample size was determined by availability; no minimum sample size for power was established. BALF samples from 1M and approximately 12M after transplantation were analyzed using MS and evaluated for intra-group (1M versus 12M) and inter-group differences (Stable vs eCLAD). All 12M samples for the eCLAD group were collected after the CLAD diagnosis.

### Study subjects

Available BALF at 1M and 12M were identified. Patients not matching the Stable or eCLAD criteria with less than five PFTs were excluded, as well as patients with biopsy-verified rejection ýA2 (16) or verified infections at time of BALF (Figure 1a). Immunosuppression, follow-up, prophylaxis protocol and clinical analyses are outlined in online supplement. All data were collected from an electronic case report form. The study was approved by the ethical review board (Dnr 791-08). All subjects provided written consent prior to inclusion. All transplants were performed in compliance with the ISHLT ethics statement.

**Figure 1.**
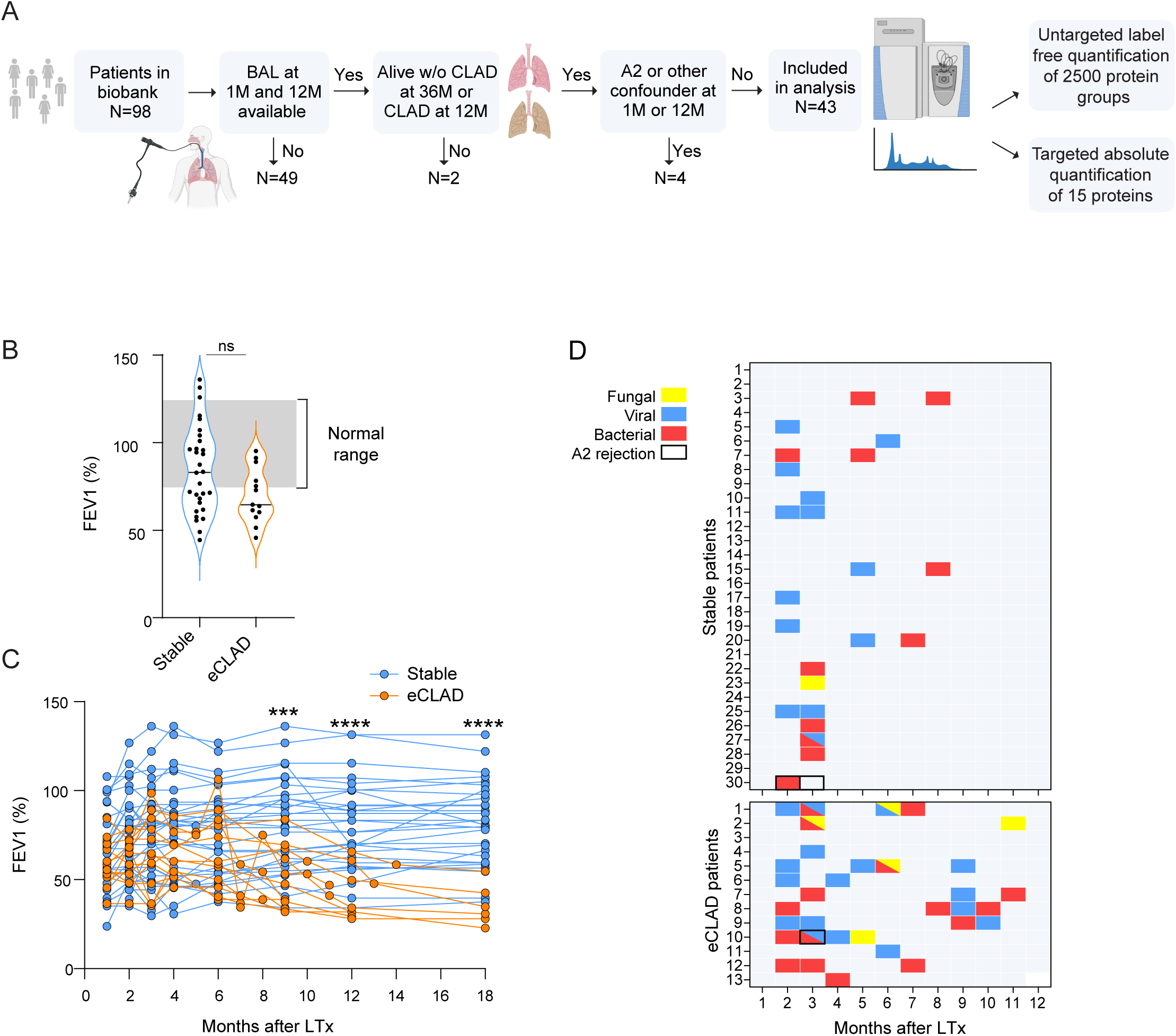
The eCLAD group of patients have respiratory infections at a later stage after lung transplantation compared to the Stable group. **A:** Selection of study subjects for mass spectrometry (MS)-based proteomics analysis of bronchoalveolar lavage (BAL) fluid. **B:** Baseline lung function (measured by % expected of forced expiratory volume in 1 second (FEV1)) of Stable and early CLAD (eCLAD) groups. Normal range indicated in grey. **C:** Lung function measurement (FEV1) over time. **D:** Respiratory infections after lung transplantation (LTx) in all Stable and eCLAD patients. Mann-Whitney U test was used for statistical analysis in B and C. ***p < 0.001, and ****p < 0.0001 for significance.

### CLAD definition

CLAD was defined according to ISHLT consensus; an irreversible 20% loss of FEV1 from the baseline in at least two consecutive PFTs at least three weeks apart, if no other reason for loss of function could be established (17). Online supplements provide further details.

### Proteomics

Samples for proteomics were prepared as previously described (12). Two types of mass-spectrometry analyses were conducted (Figure 1a): untargeted label-free analysis of all the detectable proteins and targeted absolute quantification of 15 mainly mucus or mucin-associated proteins. For targeted absolute quantification a set of heavy-labeled peptides were spiked into the samples during the sample preparation. Spiked in heavy-labeled peptides allowed to compare current results with BALF proteomics from healthy non-smoking, asymptomatic smoking and COPD patients (12).

### Proteomics data analysis

Proteomics data were analyzed with the MaxQuant program (v1.5.7.4) (18). Searches were performed against the human Uniprot protein database (downloaded 2019/12/20) and supplemented with an in-house database containing all the human and mouse mucin sequences (http://www.medkem.gu.se/mucinbiology/databases/). Proteomics data was further analyzed with Perseus software (version 1.5.5.0) (18) using default parameters. Absolute quantification of proteins that had heavy-labeled standard peptides (Table E<u>1</u>) was performed with the Skyline program (version 22.2.0.255) (19). MS proteomics data were deposited on the Proteome Xchange Consortium (http://proteomecentral.proteomexchange.org) via PRIDE partner repository (https://www.ebi.ac.uk/pride/archive/) with dataset identifier PXD044416.

### Pathway analysis

Significantly different proteins from proteomics data analysis were analyzed using Ingenuity Pathway Analysis (IPA, Qiagen).

### Gene ontology mapping

Protein-Protein Interaction Networks Functional Enrichment Analysis (STRING, https://string-db.org/) web page was used for gene ontology biological process enrichment analysis for significantly changed proteins.

### Immunofluorescence

The local pathology repository was queried for biopsies corresponding to the bronchoscopies. Available biopsies were stained for MUC5B, MUC5AC, MUC1 and CRTAC1.

### Statistical analysis

GraphPad Prism 10.4.2 (GraphPad Software, Boston, MA) was used. A p< 0.05 was considered significant. Results are presented as median ± interquartile range (IQR). For comparisons between paired measurements Wilcoxon matched pairs signed rank test was used; for comparisons between two independent measurements Mann-Whitney test was used; for three independent group comparisons, Kruskal-Wallis test with Dunn’s multiple comparisons test was used. For correlations, Spearman’s rank correlation was used. For Volcano plots of proteomic data, visualization of significantly changed proteins was done by a permutation-based false discovery rate (FDR) calculation (<FDR=0.05,S0=0.1).

## Results

### Patients and baseline clinical data

Samples from 43 patients at two time points were analyzed, 30 in the Stable group and 13 in eCLAD group (Figure 1a). The only significant difference was a higher rate of double-LTx in the Stable group (80% vs. 46%;p<0.05) (Table 1). Baseline lung function (% predicted of FEV1(20)), though variance was lower in the eCLAD group (Figure 1b). PFTs diverged significantly at 9 months (p<0.001) and in subsequent PFTs (Figure 1c). Infections and rejections grade A2+ were tracked monthly (Figure 1d). There were no significant differences during the first 3 months. Over the first year viral (69.2% vs. 33.3%;p<0.05), fungal (30.8% vs. 3.3%;p<0.05) and bacterial (69.2% vs 30%;p<0.05) infections were more frequent in the eCLAD (Table 2). However, 6/13 (46%) patients in the eCLAD group had no infections for at least 6 months before the 12M sample. Only two A2 rejections, one per group, were observed. The pattern of infections is similar early post-transplant and diverges after 2-3 months.

**Table 1.** Baseline Characteristics of Included Patients.

| Total N=43 | Stable<br>N=30(%/IQR) | eCLAD<br>N=13(%/IQR) | P Value |
| --- | --- | --- | --- |
| **Donor Age** | 49.5 (38-57.5) | 49 (39.5-62) | 0.81 |
| **Donor smoking history** | 4 (13.3%) | 1 (7.7%) | >0.99 |
| **Female Sex** | 18(60%) | 10 (77%) | 0.49 |
| **Age (years)** | 56.2(42.8-62.5) | 59.8(53.2-65.5) | 0.12 |
| **BMI** | 22.6(21.2-28.5) | 23.3(19.7-27.5) | 0.97 |
| **Smoking History** | 21 (70%) | 10 (76%) | 0.72 |
| **Double-Lung Tx** | 24 (80%) | 6 (46%) | 0.04* |
| **Diabetes** | 0 (0%) | 2 (15.4%) | 0.09 |
| **Hypertension** | 6 (20%) | 3 (23%) | >0.99 |
| **Hyperlipemia** | 2(6.7%) | 2 (15.4%) | 0.57 |
| **Transplant diagnose** |  |  |  |
| **COPD** | 9 (43%) | 7 (56.3%) | 0.72 |
| **α-1 ATD** | 4 (13.3%) | 1 (7.7%) | >0.99 |
| **Fibrosis** | 11 (36%) | 5 (39%) | >0.99 |
| **Other\*\*** | 6 (20%) | 0 | 0.15 |
| **CMV mismatch** | 6 (20%) | 2 (15%) | >0.99 |
| **Tacrolimus CNI** | 1 (3%) | 0 | >0.99 |
| **PGD 2-3** | 3 (10%) | 1 (7.7%) | >0.99 |
| **Time in ICU (days)** | 2 (1-12) | 2 (1-24) | 0.8 |
| **Baseline FEV1% predicted** | 83.1(64.7-101.8) | 64.7 (58.9-83.7) | 0.066 |

**Table 2.**
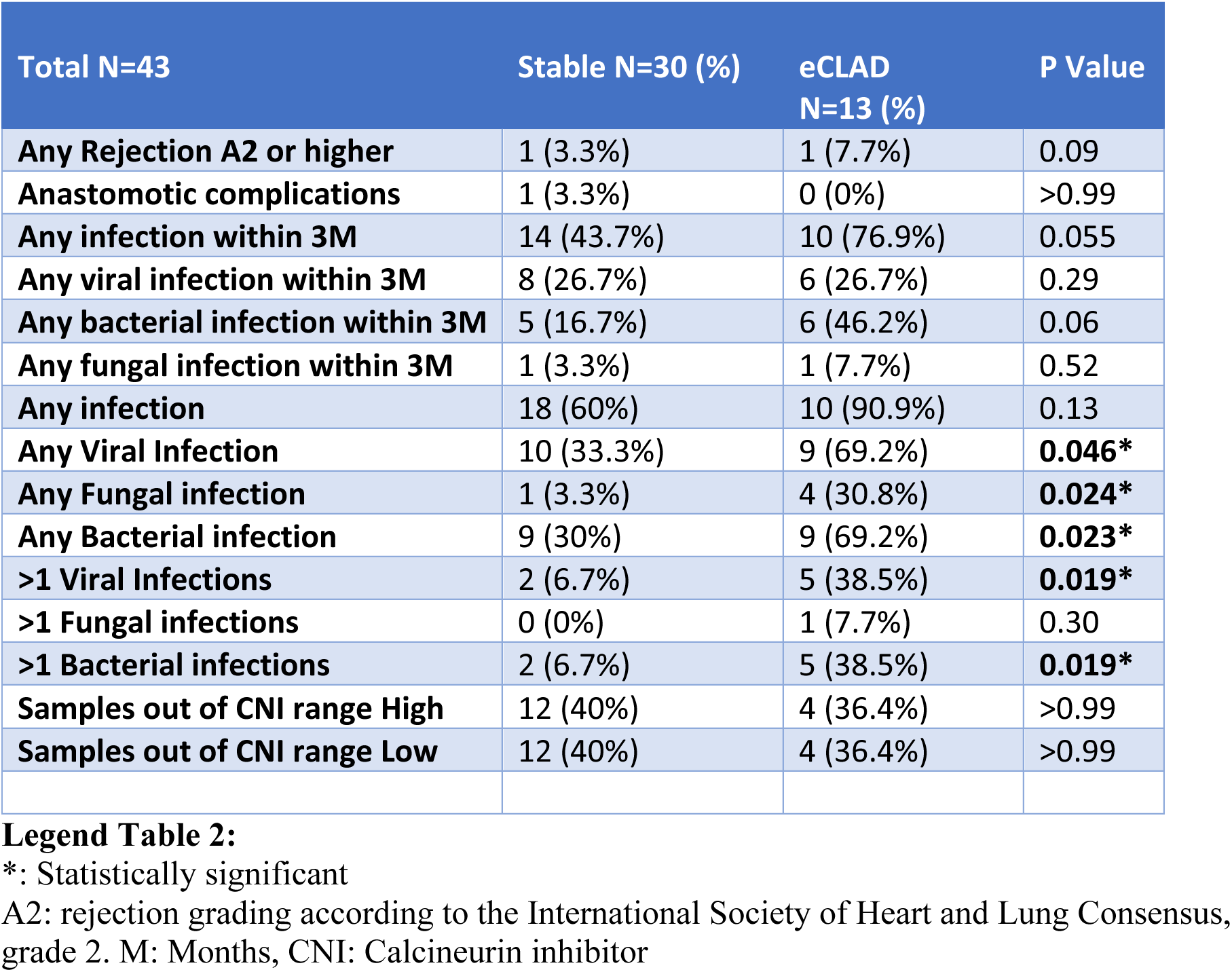
Clinical events between 1- and 12-months post-transplant.

| Total N=43 | Stable N=30 (%) | eCLAD<br>N=13 (%) | P Value |
| --- | --- | --- | --- |
| Any Rejection A2 or higher | 1 (3.3%) | 1 (7.7%) | 0.09 |
| Anastomotic complications | 1 (3.3%) | 0 (0%) | >0.99 |
| Any infection within 3M | 14 (43.7%) | 10 (76.9%) | 0.055 |
| Any viral infection within 3M | 8 (26.7%) | 6 (26.7%) | 0.29 |
| Any bacterial infection within 3M | 5 (16.7%) | 6 (46.2%) | 0.06 |
| Any fungal infection within 3M | 1 (3.3%) | 1 (7.7%) | 0.52 |
| Any infection | 18 (60%) | 10 (90.9%) | 0.13 |
| Any Viral Infection | 10 (33.3%) | 9 (69.2%) | **0.046\*** |
| Any Fungal infection | 1 (3.3%) | 4 (30.8%) | **0.024\*** |
| Any Bacterial infection | 9 (30%) | 9 (69.2%) | **0.023\*** |
| >1 Viral Infections | 2 (6.7%) | 5 (38.5%) | **0.019\*** |
| >1 Fungal infections | 0 (0%) | 1 (7.7%) | 0.30 |
| >1 Bacterial infections | 2 (6.7%) | 5 (38.5%) | **0.019\*** |
| Samples out of CNi range High | 12 (40%) | 4 (36.4%) | >0.99 |
| Samples out of CNi range Low | 12 (40%) | 4 (36.4%) | >0.99 |

### Stable patients demonstrated decreasing levels of mucins and mucus-related proteins from 1 to 12 months, whereas levels in eCLAD remain unchanged

Mucin-related proteins were quantified by targeted analysis using 56 heavy-labelled peptides for 15 proteins (Supplementary Tables E1-E2). Gel-forming airway mucins MUC5B and MUC5AC levels were ∼1 fmol/µl in both groups at 1M. Compared to healthy controls from a previous study using the same method and same set of labelled peptides (12), showed 50-100 times higher mucin levels in Stable and eCLAD patients (Figure 2a). Interestingly, MUC5B levels were similar and MUC5AC levels higher than that of COPD patients. There were no significant changes between 1M to 12M in either group for MUC5AC, MUC5B or MUC1; remaining at higher levels than seen in healthy lungs (Figure 2a). In intergroup comparisons, there were significantly more MUC5AC (p<0.01),MUC5B (p<0.01) and MUC1 (p<0.001) in the eCLAD group at 12M than in the Stable group. Immunostaining of transbronchial biopsies confirmed MUC5AC and MUC5B in both groups (Figure 2b-c). Other mucin-related proteins, significantly increased in COPD patients, including FCGBP, DMBT1, TFF3 and PSCA, showed even higher abundance in transplanted patients (Figure 2d and Figure E1b). In the Stable group, the bacterial binding protein DMBT1 (21) (p>0.01) and the goblet cell product TFF3 (p<0.05), were decreased, while the goblet cell marker FCGBP was unchanged (Figure 2d). At 12M, levels of DMBT (p<0.05) and TFF3 (p<0.01) were elevated in the eCLAD group compared to Stable (Figure 2d). These results show persistent mucin-associated abnormalities resembling COPD at 1M and 12M.

**Figure 2.**
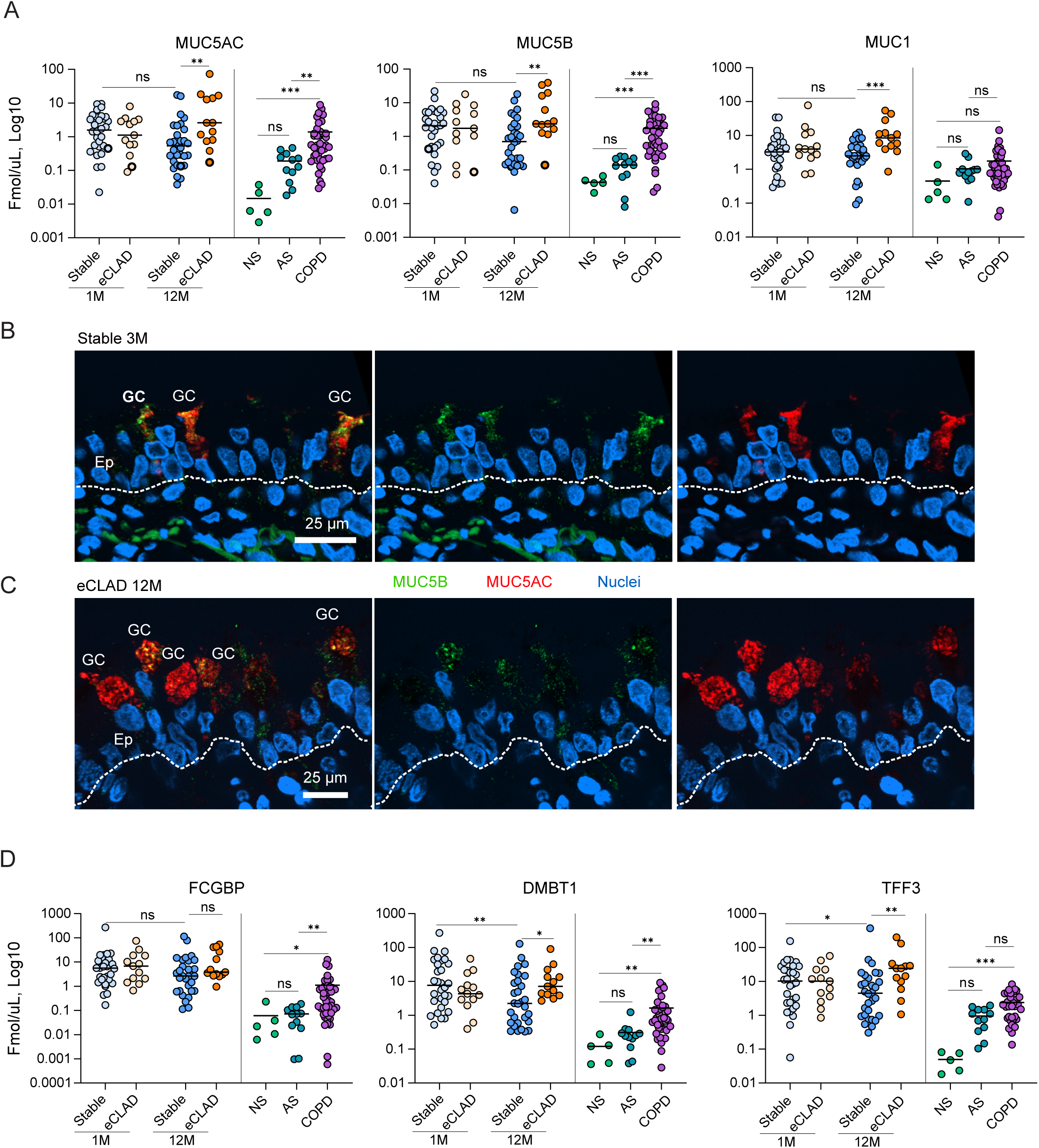
Airway mucins are increased in eCLAD patients at 12 months after lung transplantation. **A:** Absolute quantification of MUC5AC, MUC5B and MUC1 by mass spectrometry in Stable and early CLAD (eCLAD) patients at 1 month (1M) and 12 months (12M) after lung transplantation. Mucin levels are matched to a reference cohort of non-smokers (NS), asymptomatic smokers (AS) and patients with chronic obstructive pulmonary disease (COPD) from a previous study (12). Bronchial biopsies (patients marked with bold circles on A) stained by immunofluorescence to visualize mucin-producing goblet cells (GC) in a Stable patient **(B)** and an eCLAD patient **(C)**. **D:** Absolute quantification of mucin-related proteins in bronchoalveolar lavage fluid by MS from Stable and eCLAD patients at 1M and 12M compared to the reference cohort from a previous study (12). For comparisons between paired measurements Wilcoxon matched pairs signed rank test was used; for comparisons between two independent measurements Mann-Whitney test was used; for three independent group comparisons, Kruskal-Wallis test with Dunn’s multiple comparisons test was used. **p < 0.01 and ***p < 0.001 for significance.

### Stable group develops a distinct proteomic profile in BALF at 12 months

MS-based label-free (LFQ) proteomics identified 2,451 proteins, with 721 quantified after filtering proteins based on presence in a minimum of 50% of the samples in at least one group (Supplementary Table E3). The Stable group demonstrated differential levels of 155 proteins between 12M and 1M (Figure 3a, left), but no differences in BALF proteins were detected in the eCLAD group (Figure 3a, right). No intergroup differences were found at 1M (Figure 3b, left). However, 63 proteins differed significantly between groups at 12M (Figure 3b, right). Principal component analysis (PCA) confirmed no group separation at 1M, but clear separation at 12M (Figure 3c). Single vs double LTx did not influence PCA or hierarchic clustering (Figure E2a, E2b). Altogether, 195 BALF proteins were significantly changed between the 12M versus 1M Stable and 12M Stable versus 12M eCLAD, with 23 overlapping (Figure 3d). Ingenuity Pathway Analysis (IPA) of the 63 intergroup proteins at 12M revealed a “damage of lung” pathway (p=8.01E-5) (Figure 3e). IPA networks for 155 proteins in the Stable group, altered between 1M and 12M, showed reduced activity of functions at 12M, such as inflammatory cell chemotaxis, IL-8 signaling, granulocyte degranulation and killing of bacteria (Supplementary Figure E3a). Gene ontology analysis showed upregulation of proteins associated with complement activation, negative regulation of endopeptidase activity, post-translational protein modification, and platelet degranulation at 12M (Supplementary Figure E3b). It identified upregulation of proteins associated with defense response to bacteria, anti-microbial humoral response, toll-like receptor signaling pathway, and blood coagulation at 1M. In the eCLAD group, compared to the Stable group at 12M, proteins associated with defense response to bacterium, anti-microbial humoral response, toll-like receptor signaling pathway, and blood coagulation stood out (Supplementary Figure E3c). Mediators in the ”damage of lung” pathway demonstrated upregulated levels of ICAM1 and SFTPD, and downregulated levels of S100A8, AZU1, CTSG and MPO, CDL5 in Stable group at 12M (Figure 3e). These findings highlight a distinct proteomic profile in Stable group at 12M post LTx.

**Figure 3.**
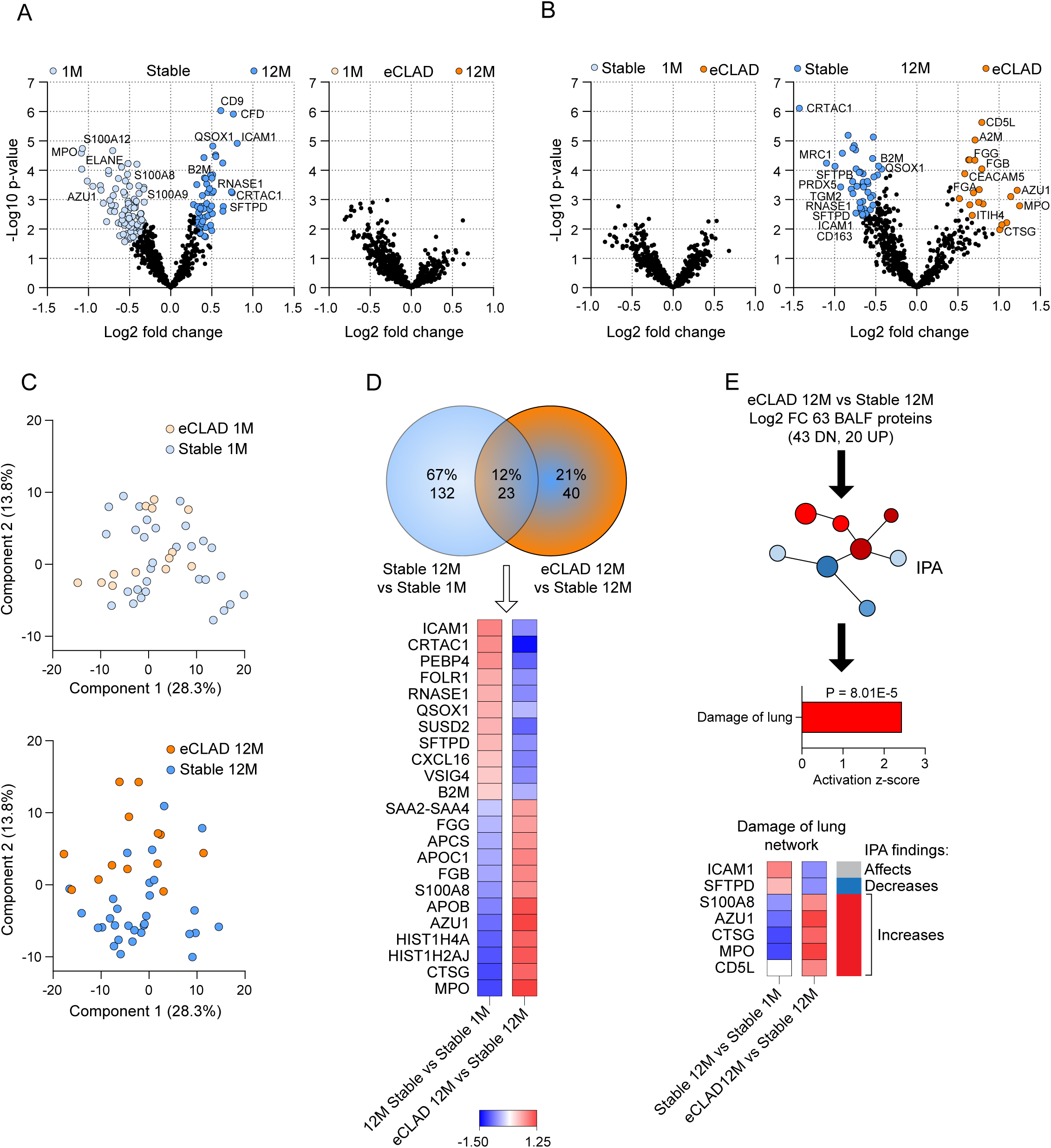
The BALF proteome changes over time in Stable patients, but not in eCLAD patients, and identifies proteins associated with lung damage. **A:** Volcano plots demonstrating differential levels of bronchoalveolar lavage fluid (BALF) proteins in Stable and eCLAD (early CLAD) groups at 1 month (1M) versus 12 months (12M) after lung transplantation. **B:** Volcano plots demonstrating differential levels of BALF proteins at 1M and 12M in Stable versus eCLAD groups. **C:** Principal component analysis of BALF proteome in Stable and eCLAD groups at 1M (above) and 12M (below). For clarity, 1M and 12M samples have been presented on separate plots. **D:** Venn diagram of significantly altered BALF proteins in Stable group at 1M versus 12M, and eCLAD group at 12M versus Stable at 12M. The 23 common proteins are shown in the heatmap. **E:** IPA analysis of all differential BALF proteins in eCLAD at 12M versus Stable at 12M identifies damage of lung network. Key proteins are shown in the heatmap with actual measurement (left) and IPA predicted direction for lung damage (right). The heatmap key (D) corresponds to Log2 ratios for D and E.

### Lower neutrophil and higher macrophage frequency in the Stable group at 12 months

Differential cell-count in BALF showed higher levels of neutrophils at 12M in the eCLAD group compared to Stable group (p < 0.05) (Figure 4a). Macrophages were lower in the eCLAD group at 12M compared to Stable group at 12M (p < 0.01) (Figure 4a). Protein levels determined by MS in BALF reflected cellular infiltrates, where macrophage frequency demonstrated a positive correlation with the macrophage product MRC1 (Figure 4b, left) and neutrophil frequency correlated positively with the neutrophil product MPO (Figure 4b, right). The abundance of mediators identified in the “damage of lung” pathway (ICAM1, SFTPD, A100A8, AZU1, CTSG, MPO) correlated with change in lung function in the Stable and eCLAD groups at 12M (Figure 4c). These findings further support that the changes found in the Stable group are related to a beneficial outcome after LTx.

**Figure 4.**
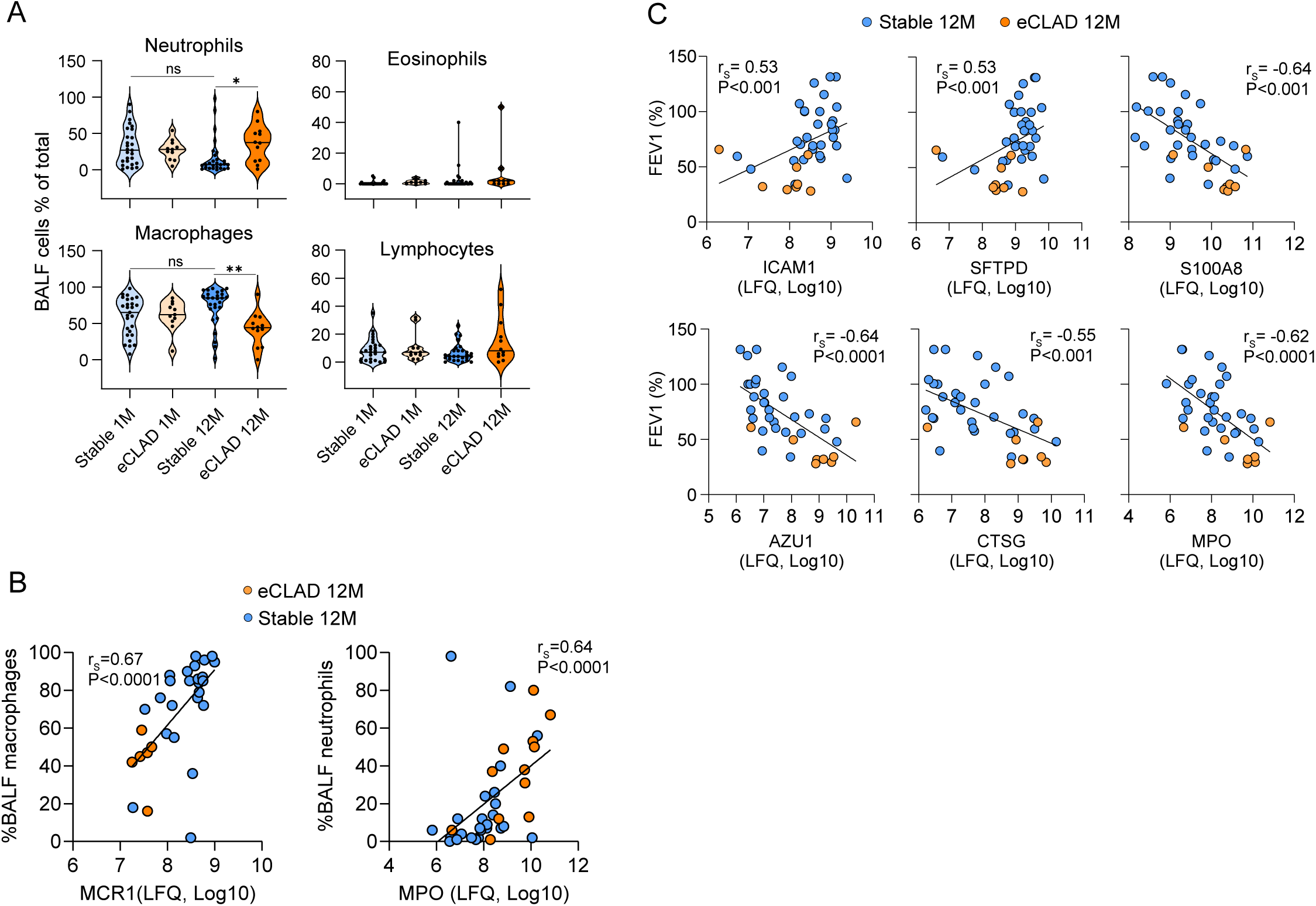
Lung damage-markers correlate with improved lung function in the Stable group. **A:** Differential cell count of bronchoalveolar lavage fluid (BALF) cells at 1 month (1M) and 12-months (12M) after lung transplantation in Stable and early CLAD (eCLAD) patients. **B:** Spearman correlation between BALF macrophage frequency and level of macrophage-associated protein MCR1 in BALF (left), and BALF neutrophil frequency and level of neutrophil-associated protein MPO in BALF (right). **C:** Spearman correlation between lung damage-associated proteins in BALF and lung function measurement by % expected forced expiratory volume in 1 second (FEV1). For comparisons between paired measurements Wilcoxon matched pairs signed rank test was used; for comparisons between two independent measurements Mann-Whitney test was used in A. *p < 0.05, and **p < 0.01 for significance.

### CRTAC1 levels in BALF correlelate to lung function after lung transplantation

CTRAC1, previously linked to healthier lungs (21), was significantly higher at 12M in the Stable group compared to 1M (p>0.05) and significantly less abundant in the eCLAD group than in Stable group at 12M (p<0.01) (Figure 5a). At 12M, CRTAC1 levels in Stable patients were closer to those in healthy individuals than asymptomatic smokers or COPD patients (12) (Figure 5a). Biopsies showed less CRTAC1 staining in 1M eCLAD than in the 1M Stable (Figure 5b) despite no significant difference in BALF protein abundance at that time (Figure 5a). CRTAC1 intensity remained weak in the 12M eCLAD biopsies (Figure 5c), with lower signal intensity than in the 1M Stable (Figure 5b). No 12M stable biopsies were available for comparison. Levels of CRTAC1 correlated directly with lung function in the Stable and eCLAD groups at 12M but not at 1M (Figure 5d). Furthermore, the abundance of CRTAC1 in BALF correlated with all the significantly changed proteins annotated as belonging to the “damage of lung” pathway (ICAM1, SFTPD, A100A8, AZU1, CTSG, MPO) (Figure 5e). These findings suggest that the sensitivity in CRTAC1 levels in BALF could help separate Stable patients from eCLAD patients.

**Figure 5.**
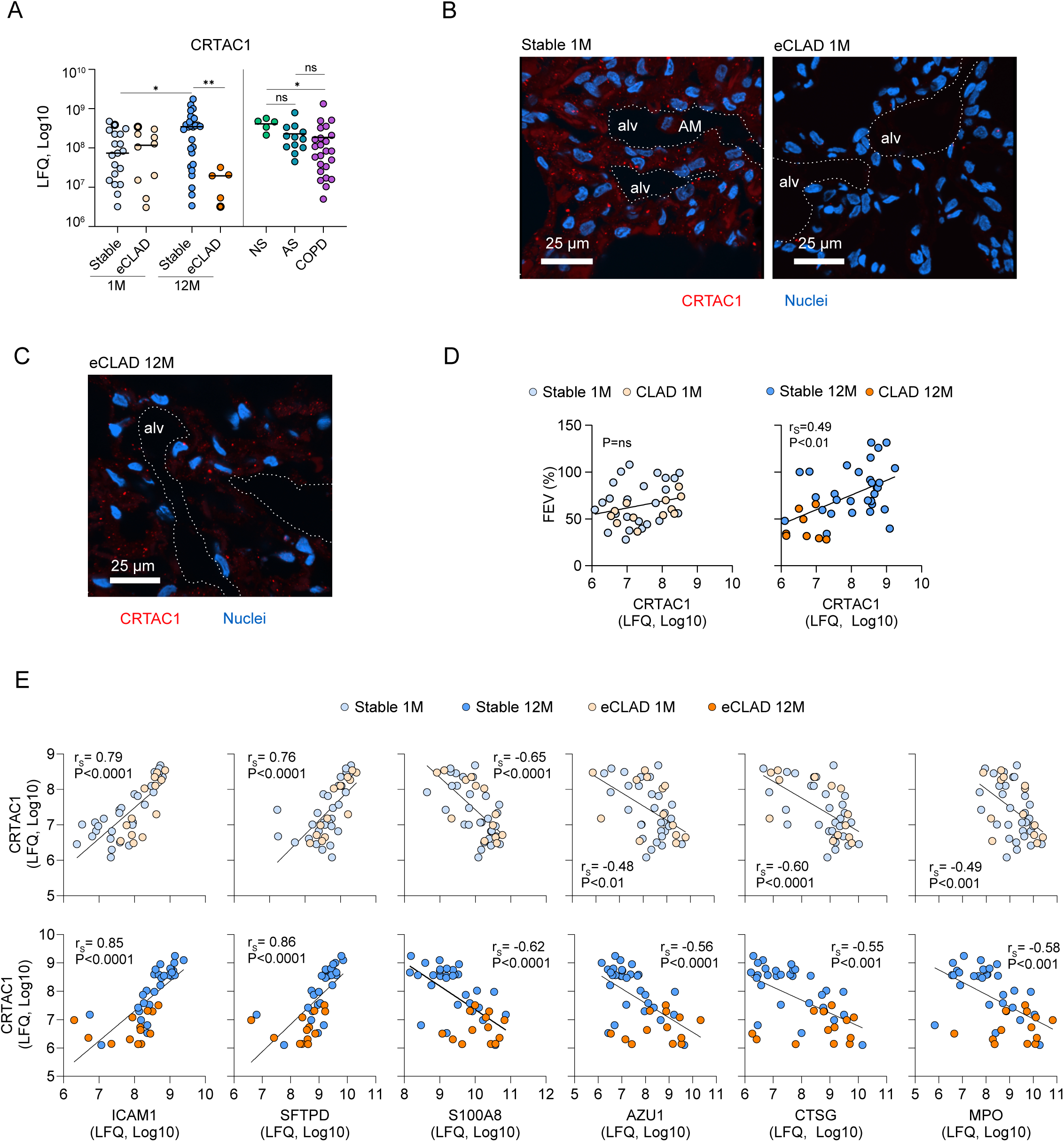
CRTAC1 levels are increased in Stable patients at 12 months after lung transplantation and correlate with lung function measurements. **A:** CRTAC1 levels in bronchoalveolar lavage fluid (BALF) determined by mass spectrometry in Stable and early CLAD (eCLAD) patients at 1 month (1M) and 12 months (12M) after lung transplantation, compared to a reference cohort of non-smokers (NS), asymptomatic smokers (AS) and patients with chronic obstructive pulmonary disease (COPD) from a previous study (12). CRTAC1 stained by immunofluorescence in bronchial biopsies obtained at 1M **(B)** and 12M **(C)** (patients marked with bold circles on A). **D:** Spearman correlation between CRTAC1 levels in BALF and lung function measurement by % expected forced expiratory volume in 1 second (FEV1). **E:** Spearman correlation between CRTAC1 levels in BALF and levels of lung damage-associated proteins in BALF. For comparisons between paired measurements Wilcoxon matched pairs signed rank test was used; for comparisons between two independent measurements Mann-Whitney test was used; for three independent group comparisons, Kruskal-Wallis test with Dunn’s multiple comparisons test was used on A. *p < 0.05, **p < 0.01 for significance

## Discussion

The biological mechanisms of CLAD (17) and successful lung adaptation post-transplant remains poorly understood. In this study, the Stable and the eCLAD patients showed similar BALF protein profiles at 1M after transplantation but began to diverge after three months post-LTx which possibly reflects early mucosal defense stabilization. By 12M, the Stable group showed BALF proteome normalization, unlike the eCLAD group, which was unchanged between timepoints. This suggests more effective lung adaptation in the Stable group while the eCLAD group do not progress and adapt for unknown reasons.

Excluding major confounders, such as infections and grade ≥2 rejection at sampling, strengthens the link between BALF proteome changes and the underlying pathophysiological mechanisms. The high number of single lung transplants (SLTx) reflects common practice at the time (22). SLTx is a risk factor for shorter CLAD-free survival without differences in the number of infections (23). No significant difference in best PFT and the ≥20% decline in the eCLAD group suggests intrinsic allograft issues rather than procedural differences. This is supported by the lack of distinction between SLTx and double LTx in BALF proteome.

Due to study design, we cannot conclude whether the higher number of infections in the eCLAD group impedes adaptation or result from it. However, infection rates did not differ during the first three months. There is no difference in immunosuppression nor missed through-calcineurin inhibitor levels, which excludes treatment differences. The BALF cell reflects known CLAD patterns with more neutrophils than macrophages (24). The observed period (1M-12M) overlaps with delayed mucociliary clearance (13,14). Retarded mucus clearance is a protection mechanism that limits bacteria reaching the epithelial cells (12), suggesting that mucus system dysfunction may contribute to the infection pattern.

Lower MUC5AC, MUC5B and MUC1 levels in the Stable group at 12M support a role for mucus clearance in eCLAD development. However, both groups showed mucin levels more like the COPD than non-COPD controls from our previous study using the same method (12). This indicates ongoing stress even in the Stable allografts. The impact of pre- and post-operative ventilation on 1M is unclear, though MUC5AC is linked to ventilator-induced lung injury (25). High MUC5B and MUC5AC mucins in biopsies from both groups further support that the transplanted lungs differ from healthy controls (12). Previous studies, also report elevated mucin levels in BALF from CLAD patients with high MUC5B levels associated with shorter post-diagnosis survival (26,27).

Overall, the protein levels of the 12M Stable group stood out compared to the other groups (1M Stable, 1M eCLAD and 12M eCLAD). The network identified through IPA contained proteins associated with acute lung injury (MPO, CTSG, ICAM1) (28), neutrophil activity (MPO, CTSG, S100A8) (28), and emphysema (SFPTD) (29). In support of neutrophil activity identified in the network, S100A8/A9 proteins have been associated with CLAD (30), MPO is associated with neutrophil extracellular traps (31) and ELANE, which is also elevated, is central in the process of forming these (32). Interestingly, all these proteins have been identified as possible therapeutic targets (34–37). The direct correlation between protein levels in BALF and pulmonary function is a further indication of the central role of ICAM1, SFPTD, S100A8, AZU1, CTSG and MPO in post-transplant lung adaptation and is interesting in the context of recent observations highlighting the importance of best post operative-lung function (38). The finding also mechanistically supports early post-operative introduction of Azithromycin for neutrophil modulation (39).

CRTAC1, a glycosylated extracellular matrix protein originally found in human cartilage (40) but also produced by type 2 alveolar cells (41), was increased over time in the Stable group. While its function in the lung is still unknown, CRTAC1 has been negatively associated with lung fibrosis (21). In this study it is strongly correlated with the “damage of lung” network and lung function for both the eCLAD and the Stable groups. This suggests a central mechanistic role in lung health, and a potential for further exploration as a biomarker.

This study has limitations. Although based on a prospective cohort, the retrospective design may introduce hidden bias and preclude mechanistic conclusions. Most patients received cyclosporin, now considered a suboptimal treatment option after lung transplantation (42), and the small number of CLAD patients limits phenotype-specific analysis. Strengths of the study are the very well-defined infection-surveilled prospective cohort and the clear definition of confounder-depleted grouping. Furthermore, the general protocolized surveillance reduces risk of group-dependent data collection bias.

In conclusion, this study suggests that stable, long-term surviving allografts undergo an adaptation towards a healthier state. However, the mucin proteomic profile, affecting first-line infection defense, remains like that of COPD lungs, even in clinically stable allografts. This suggests that even allografts with excellent function and CLAD-free survival are under functional stress corresponding to a chronic lung disease. We found that the functional status of the transplanted lung can be directly correlated to the proteins in the identified lung damage network. Notably, CRTAC1 correlates with all these processes, making it an interesting protein in association with pulmonary stability.

Our findings reveal a gap between the transplanted lung and healthy lungs, underscoring the need for further improvement of allograft management pre-, peri-, and post-operatively to enhance transplant outcomes. The correlation of functional status with closely interacting proteins in the damage of lung network suggests approaches for mechanistic and therapeutic research to be validated in future prospective studies. While reversing CLAD remains a major goal for the lung transplant community, the current study supports that research aiming to explore, improve and preserve best initial lung function should be a parallel priority to advance lung transplantation outcomes.

## Abbreviations

LTx: Lungtransplantation
CLAD: Chronic Lung Allograft Dysfunction
COPD: Chronic obstructive lung disease
eCLAD: Early CLAD (group denomination)
1M: One Month
12M: 12 Months
BALF: Bronchoalveolar Lavage Fluid
PFT: Pulmonary Function Test
FEV1: Forced expiratory volume during the first second
FDR: False discovery rate

## Acknowledgements

The following sources funded the study: The project has been supported by the Gothenburg medical society (GLS-986352) Knut and Alice Wallenberg Foundation (2017.0028), Swedish Research Council (2023–02474), the European Research Council ERC (101100663, 694181), IngaBritt and Arne Lundberg Foundation (2018-0117), Västra Götalandsregionen (ALFGBG-440741, -236501, -995932, -942865(JMM), Wilhelm and Martina Lundgren’s Foundation.

The Swedish Heart-Lung Foundation (20190311, 20200680, 20220404 (AE); 20210377, 20230413 (GCH) +JMM); 20220466 JMM), Lederhausen’s Center for CF Research at Univ. Gothenburg, the Wenner-Gren Foundations (KJ) and Herman Krefting Foundation (KJ)), Sahlgrenska, NHV -967493.

The following disclosures are reported: LA,KJ - no disclosures, AE – Shareholder Mucolife Therpeutics, MG – Speakers fee, Therakos (UK) DSMB E-CLAD study, TP – Shareholder and CEO Mucolife Theraputics, – Mucolife Therapeutics, JW - No disclosures, GCH – Shareholder and CSO, Mucolife Therapeutics, JMM – Speakers Fee Boehringer Ingelheim, AstraZeneca, Takada Pharma, GlaxoSmithKline, Mallinckrodt, Vicore Pharma; Advisory Board - Boehringer Ingelheim; Boardmember (unpaid) Swedish pulmonary fibrosis registry, Scandinavian Heart and Lung group.

In addition to this we wish to acknowledge the work of the staff of the lung transplant department and the staff of the viral detection laboratory at Sahlgrenska University for their support of the current study.

---

**Figure E1.**
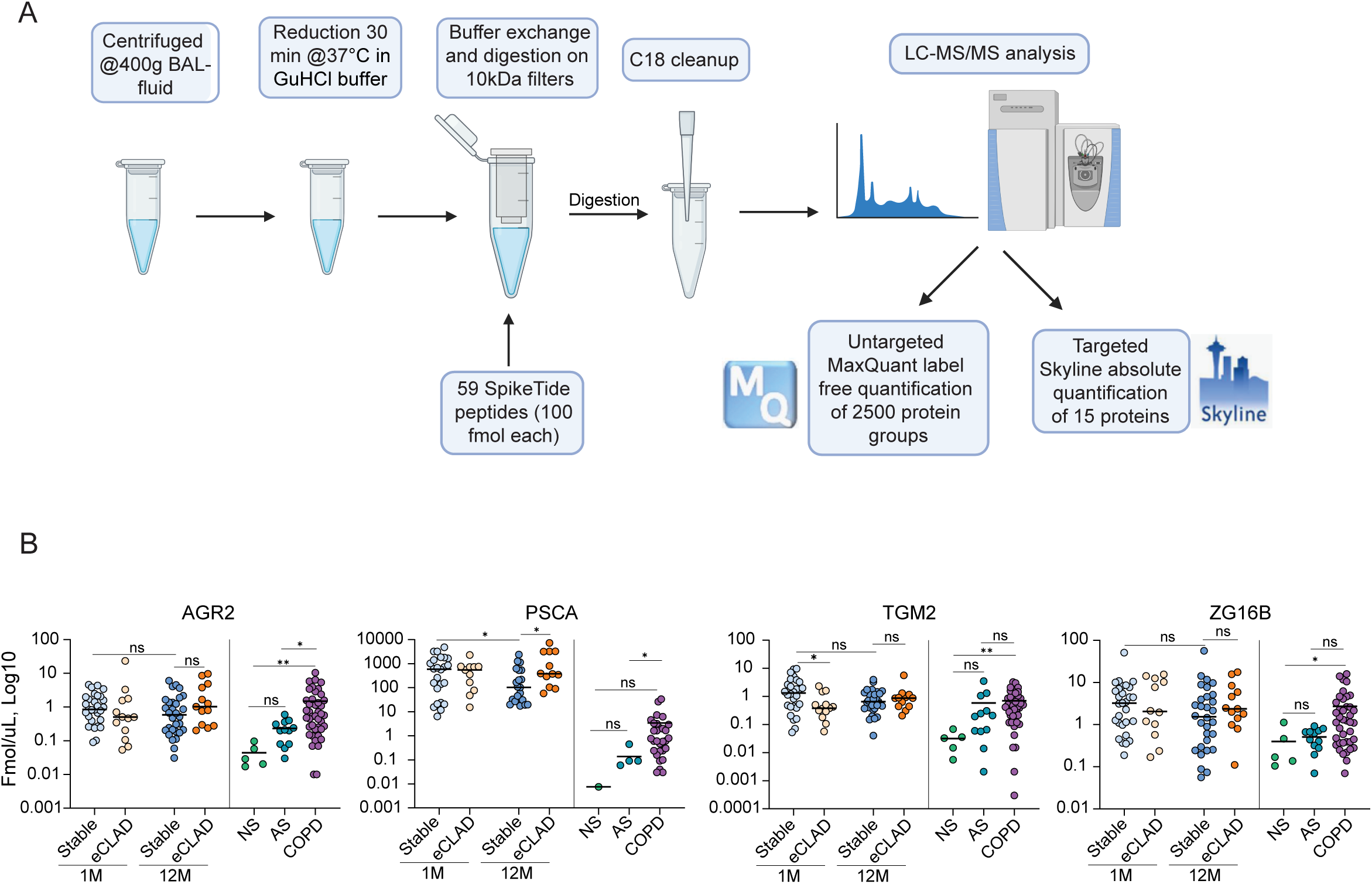

**Figure E2.**
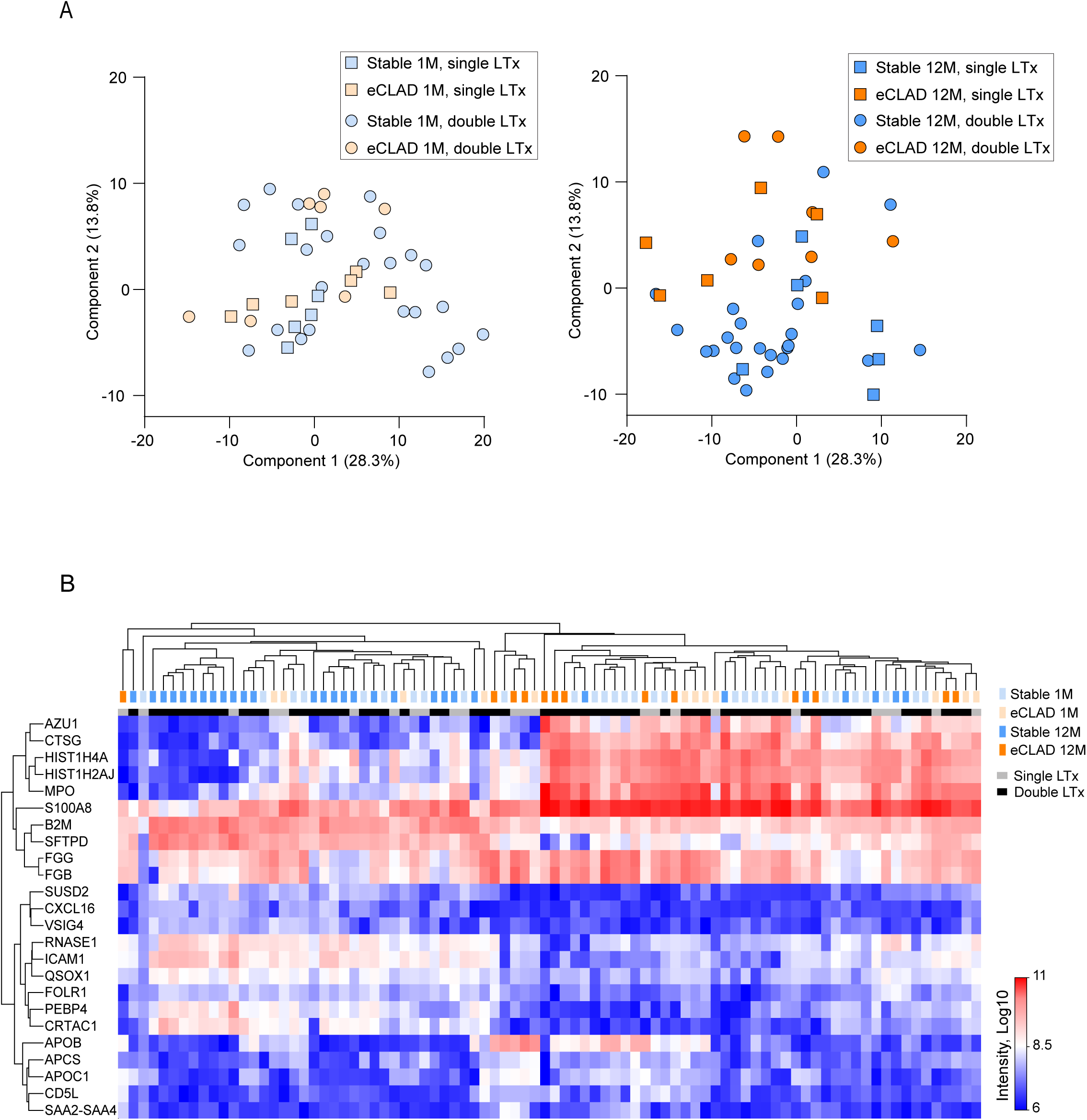

**Figure E3.**
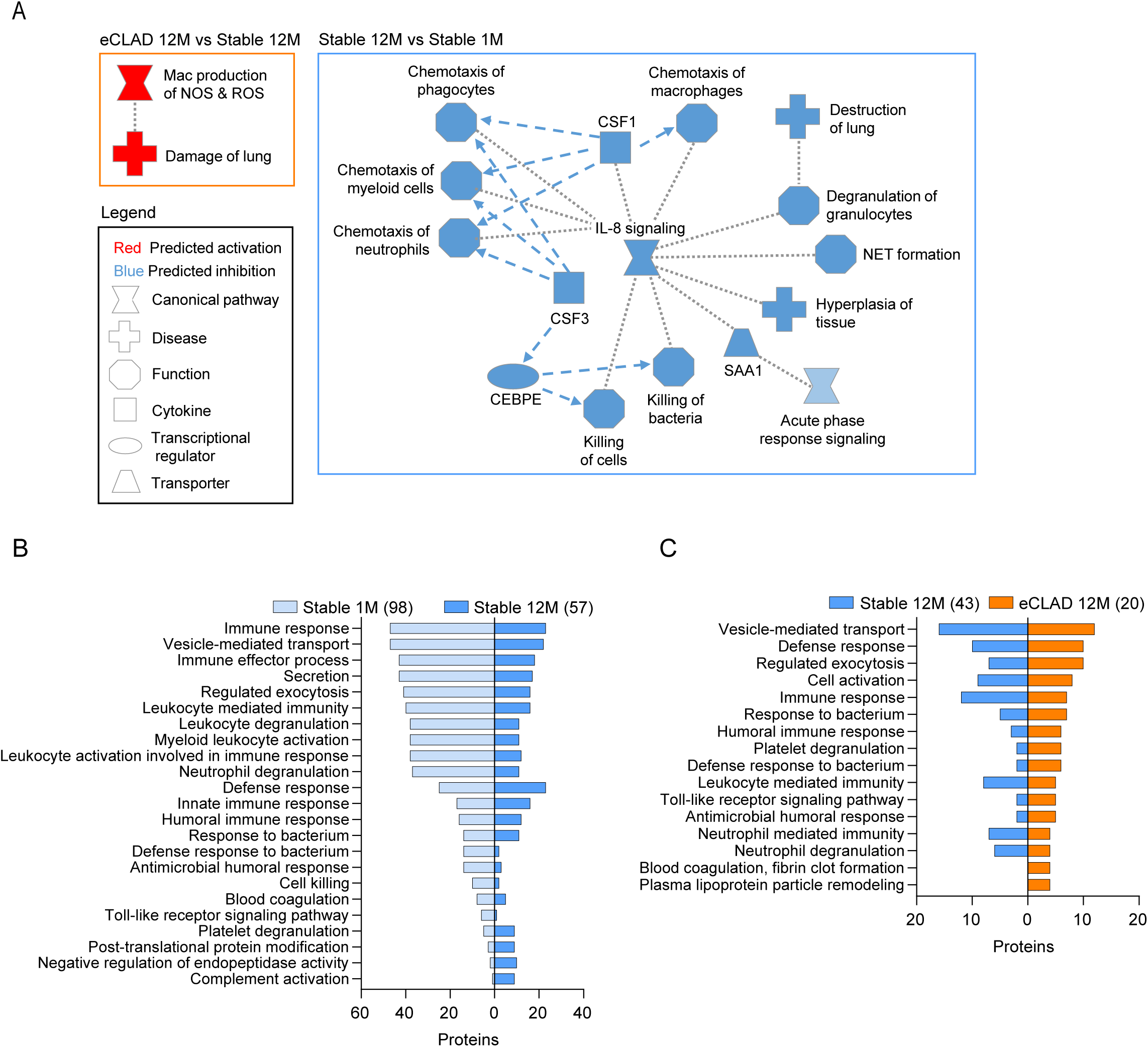

**Figure E4.**
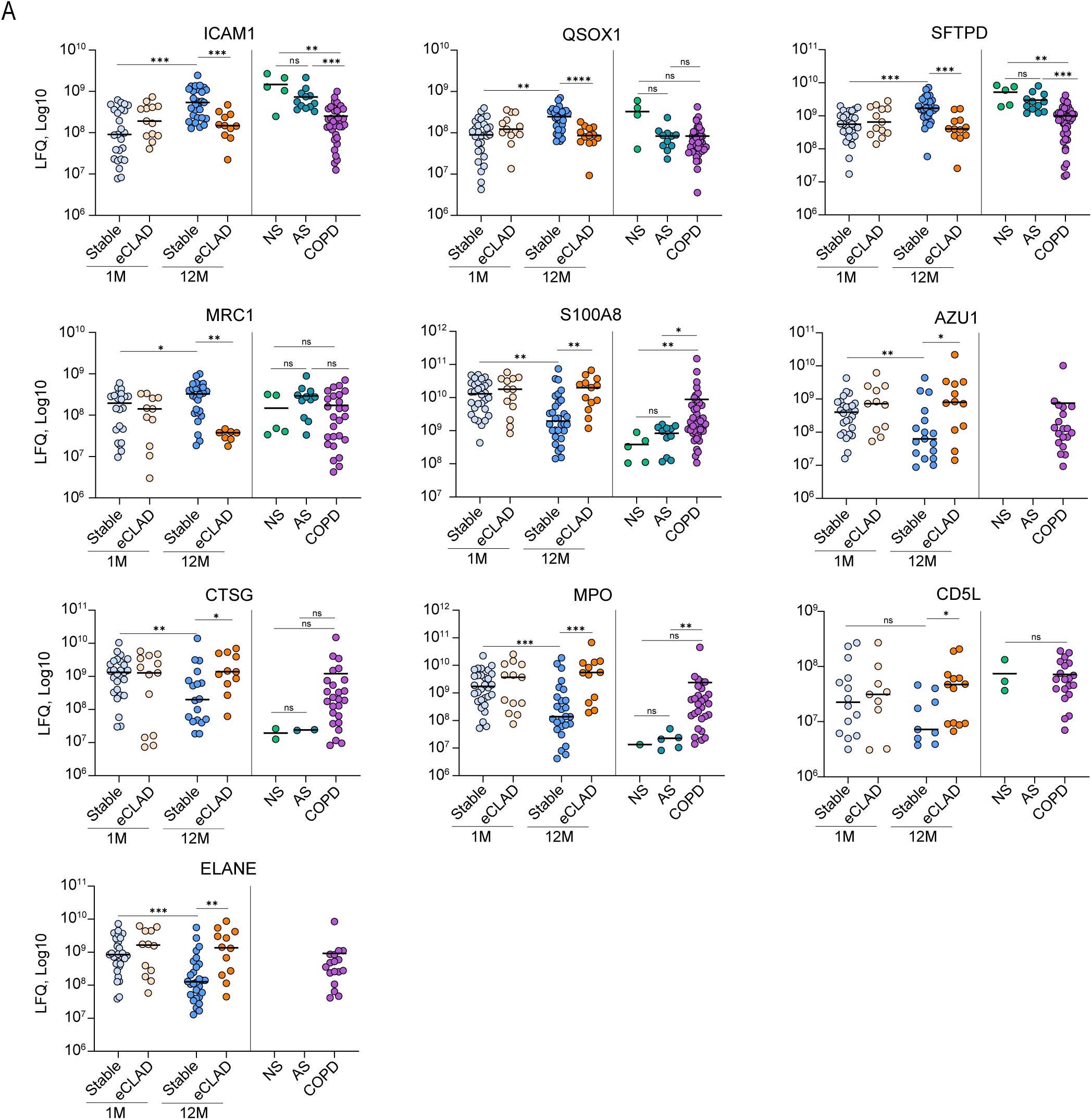

