## Supplemental information for "Proteomic analysis of bronchoalveolar lavage fluid after lung transplantation associates stable allograft function with less lung damage at 12 months"

Online data supplement

Supplemental methods.

**Immunoupression and prophylaxis**

Induction therapy consisted of rabbit antithymocyte globulin which was given for 1-3 consecutive days together with methylprednisolone IV. Post-transplantation immunosuppression included prednisone, 0.3 mg/kg/day and mycophenolate mofetil (MMF), 2 g/day. The patients then received either oral Cyclosporine (CSA) (1-2 mg/kg) adjusted to maintain a serum level of 300-350 ng/mL or Tacrolimus (TAC), 0.075 mg/kg given orally divided into two doses daily adjusted to maintain a serum level of 14-16 ng/mL. The dosage of immunosuppression was gradually lowered during follow-up. Further changes in immunosuppressive therapy were based on clinical presentation.

All patients initially received a combination of 80 mg of Sulfamethoxazole and 400 mg of Trimethoprim thrice weekly. Patients who tested positive for cytomegalovirus (CMV) antibodies at the pre-transplant evaluation were given 900 mg Valganciclovir daily for 3 months; the rest of the patients were given the same treatment for 6 months. Adjustments were made based on clinical response. All patients were treated with simvastatin 10 mg once daily.

**Follow-up protocol, BALF and clinical controls**

***Follow-up protocol***

Outpatient visits were performed at 1, 2, 3, 4, 5, 6, 9, 12, 18, and 24 months and then every year up until five years. Patients self-monitored with home spirometry and a persistent drop in FEV1 of 10% also prompted an extra visit as did symptoms of infection. At every visit, a Pulmonary function test (PFT) and chest x-ray were performed, patients were tested for CMV and Eppstein-Barr virus as well as torque-teno virus and a nasopharyngeal swab for viral detection were performed. Mandatory bronchoscopies were performed at 1, 3 and 12 months after transplantation. For-cause bronchoscopies were performed at extra visits prompted as previously described. BALF was collected and cultured in every bronchoscopy for fungal, bacterial and mycobacteria infection. Furthermore, the BALF was analyzed for viral infections according to the header respiratory viral analysis, CMV, *Legionella pneumophila* and *Pneumocystis jirovecii* using PCR. Computed tomography was performed for cause.

***BALF collection***

BALF samples were collected according to a standardized protocol. After a brief lung inspection, the BALF was collected by infusion of 3 x 50 mL 37°C sterile pyrogen-free phosphate-buffered saline aliquots into a segmental middle lobe or lingula bronchus, with the bronchoscope in a wedged position. A 60-80% return of instilled fluid was considered acceptable. Part of the fluid was transferred to a separate container in a sterile manner, placed on ice, and then frozen at -80° C until analysis.

***Differential leukocyte count***

All BALF samples underwent a differential leukocyte count. After cytospin, the BALF cells were stained with May-Grünwald Giemsa stain, and 200-500 cells were counted and classified according to leukocyte type.

***Respiratory viral analysis***

For viral detection both in nasopharyngeal swabs and BALF, a multiplex PCR in-house method was used with the ability to detect adenovirus, bocavirus, *Chlamydophila pneumoniae*, human coronaviruses (CoV) NL63, HKU1, OC43 and 229E, human enterovirus, human metapneumovirus, human rhinovirus (hRV), influenza A and B, *Mycoplasma pneumoniae*, parainfluenza virus (1, 2 and 3) and respiratory syncytial virus.

***Chronic lung allograft dysfunction (CLAD) definition***

CLAD was defined as an irreversible 20% loss of forced expiratory volume during the first second (FEV1) from the baseline in at least two consecutive PFTs at least three weeks apart if no other reason for loss of function could be established after actively searching for confounders such as infection, bronchial stenosis, pleural effusion among others. Baseline FEV1 was the mean of the two best post-transplant FEV1 measurements. The time for CLAD diagnosis was defined as the time of the first consecutive PFT used to establish the condition. The CLAD diagnosis was verified independently by two experienced LTx specialists. At the time of the study, only the ISHLT consensus statement defined only the BOS phenotype and overlapped with the current CLAD definition. The RAS phenotype, as proposed by Sato et al. in 2011, existed, but the RAS consensus paper was not published. The mixed and undefined phenotypes were not present. Furthermore, several publications from the Toronto group suggest a fifth phenotype currently not outlined in the consensus document. This was published post  The study uses the CLAD umbrella term for the main analysis. Phenotyping was performed for exploratory reasons. The original blinded double validation of the diagnosis remained unchanged in the current study; all phenotyping was performed according to current guidelines as briefly summarized in the following table. Obstruction was defined as a FEV1/FVC quota below 0.7. Restriction was defined as a total lung capacity (TLC) drop of 10% or more compared to baseline TLC defined as the TLC measurement obtained at the same time as or near the best two post-operative FEV1 measurements.

| Phenotype | Obstruction | Restriction | Radiological Opacities |
|---|---|---|---|
| **BOS** | Yes | No | No |
| **Mixed** | Yes | Yes | Yes |
| **RAS** | No | Yes | Yes |
| **Undefined** | Yes | No | Yes |
| **Undefined** | Yes | Yes | No |
| **Unclassifiable** | No | No | No |
| **Unclassifiable** | No | No | Yes |
| **Unclassifiable** | No | Yes | No |

Legend BOS: Bronchiloitis obliterans Syndrome, RAS: Restrictive Allograft Syndrome

### Detailed Mass spectrometry description

#### *Sample preparation for mass spectrometry*

BALF samples (90 µl) were centrifuged for 10 min, 400xg, at 4°C. Supernatants (80 µl) were collected and mixed with 80 µl of reduction buffer (6 M GuHCl, 0.1 M Tris/HCl pH 8.5, 0.1M DTT, 5 mM EDTA). Samples were reduced by mixing 30 min at 800 rpm, 37°C. An additional 50 µl of 6 M GuHCl was added and samples were transferred to 10 kDa cut-off filters (Nanosep centrifugal devices with OMEGA filter, PALL, Port Washington, NY). Proteins were alkylated and subsequently digested for 4 hours with LysC (Wako Chemicals), followed by overnight trypsin (Promega) digestion. Heavy peptides (JPT Peptide Technologies) for absolute quantification (59 peptides for 15 proteins, 100 fmol each, listed in Table E1) were added before trypsin digestion. Peptides were released from the filter by centrifugation and cleaned with in-house made StageTip C18 columns prior to MS analysis (Figure E1a).

MS analysis was performed on an EASY nanoLC 1200 system (Thermo Fisher Scientific), connected to a Q-Exactive HFx hybrid quadrupole-Orbitrap mass spectrometer (Thermo Fisher Scientific) through a nanoelectrospray ion source. Peptides were separated in an in-house packed C18 reverse-phase column (150 × 0.075 mm inner diameter, C18-AQ 3 µm) by a 35-minute gradient from 5%–30% B, followed by 30%-45% B in 5 min (A, 0.1% formic acid; B, 0.1% formic acid/80% acetonitrile 250 nl/min). Full mass spectra were acquired from m/z 350–1,600, with a resolution of 60,000. Up to 15 of the most intense peaks (charge state ≥2) were fragmented with a normal collision energy of 27%. Tandem mass spectra were acquired with a resolution of 15,000 and dynamic exclusion of 20 s. For absolute quantification, a separate targeted MS method (parallel reaction monitoring) was used, where only precursors and fragments of the heavy and corresponding light peptides were scanned with a resolution of 15,000.

### Detailed Proteomics data analysis

Proteomics data were analyzed with the MaxQuant program (version 1.5.7.4) Searches were performed against the human Uniprot protein database (downloaded 2019/12/20) and supplemented with an in-house database containing all the human and mouse mucin sequences (http://www.medkem.gu.se/mucinbiology/databases/). Full tryptic specificity was used, with a maximum of 2 missed cleavages, a precursor tolerance was set to 20 ppm in the first search followed by 7 ppm for the main search, and 0.5 Da for fragment ions. Carbamidomethylation of cysteine was set as a fixed modification, and methionine oxidation as well as protein N-terminal acetylation were set as variable modifications. The false discovery rate (FDR) was set to 1% both at the peptide and protein levels, and the minimum required peptide length was set to 6 amino acids. Proteins were quantified by the MaxQuant label-free quantification (LFQ) algorithm, using a minimum of 2 peptides. Data filtering, clustering, principal component analysis and Volcano plots were performed with the Perseus software (version 1.5.5.0) (18) using default parameters. Reverse hits and common contaminants were removed, highly abundant albumin was excluded as a contaminant. The

data was filtered based on the presence of a protein in at least 50% of the samples in one of the groups (1M Stable, 12M Stable, 1M eCLAD and 12M eCLAD). Missing values were replaced with the normal distribution of values. Significantly different proteins were determined with a permutation-based FDR calculation (FDR=0.05, S0=0.1).
Absolute quantification of proteins that had heavy-labeled standard peptides (Table E1) was performed with the Skyline program (version 22.2.0.255) (19). The average peptide concentration was reported as protein concentration. The MS proteomics data were deposited on the Proteome Xchange Consortium (http://proteomecentral.proteomexchange.org) via the PRIDE partner repository (https://www.ebi.ac.uk/pride/archive/) with the dataset identifier PXD044416.

**Immunoflourecence**

MUC1: Biopsies from human airway epithelium were fixed in 4% paraformaldehyde, embedded in paraffin, sectioned onto glass slides and baked for 2 hours at 60°C. Slides were deparaffinized in xylene substitute (2 × 10 min, 60°C), and rehydrated in 100% ethanol (10 min), 70% (v/v) ethanol (5 min), 50% (v/v) ethanol (5 min), and 30% (v/v) ethanol (5 min). Slides were placed in antigen retrieval buffer (0.01M citric acid, pH 6.0) at 100 degrees for 10 min and then cooled to RT (2 h) and transferred to PBS. Tissue sections were enclosed with a hydrophobic PAP pen, permeabilized with 0.1% Triton X-100 in PBS for 5 min and blocked with 5% fetal calf serum (FCS) in PBS for 1 h at room temperature (RT). Primary (overnight at 4°C) and secondary antibodies (2h at RT in the dark) were added in dilution buffer (5% FCS in PBS). Primary antibodies were rabbit anti-MUC1 CT1 polyclonal antibody (1:200) (1) and mouse anti-tubulin (acetylated) monoclonal antibody (1:1000) (Sigma-Aldrich Cat# T6793, RRID:AB_477585). Secondary antibodies were donkey anti-rabbit IgG Alexa Fluor 488 (1:1000) (Thermo Fisher Scientific Cat# A-21206 (also A21206), RRID:AB_2535792) and goat anti-mouse IgG Alexa Fluor 647 (1:1000) (Thermo Fisher Scientific Cat# A-11001, RRID:AB_2534069). Nuclear DNA was stained with 5 µg/mL Hoechst 34580 (ThermoFisher Scientific Cat# H21486) for 10 min. Slides were washed three times with PBS after each incubation step. Coverslips were mounted using Prolong Gold antifade (ThermoFisher Scientific Cat# P36980) and polymerized for 2 h at RT. Slides were imaged using an upright LSM 700 Axio Examiner Z.1 confocal imaging system (Carl Zeiss).

MUC5AC, MUC5B and CRTAC1: Transbronchial biopsies were fixed in 4% formalin, embedded in paraffin and cut in 4 µm sections according to clinical routine procedures. Sections were dewaxed in Xylene (Sigma, St. Louis, MO) and rehydrated. Antigen retrieval was performed by heating in 100˚C for 20 min and then cooling at room temperature for 20 min in 0.01 M citric buffer pH 6 for MUC5AC and MUC5B or 10 mM Tris-HCl with 1 mM EDTA at 8.0 for CRTAC1. Sections were blocked for 60 min with 3% donkey serum in Tris-buffered saline (TBS) and permeabilized with 0.1% Triton X-l00. Stainings for MUC5B and MUC5AC were performed sequentially. Primary antibodies, mouse monoclonal anti-MUC5AC clone 45M1 (Cat# MA1-38224, RRID:AB_2146842, ThermoFisher Scientific, Waltham, MA), rabbit polyclonal anti-MUC5B, made in house (Fakih et al., 2020 doi: 10.1152/ajplung.00485.2019), or rabbit polyclonal anti-CRTAC1 (Cat# PA5-78519, RRID:AB_2735688, Invitrogen, ThermoFisher Scientific, Waltham, MA), were incubated in blocking solution over night at 4°C. For detection of MUC5B, donkey anti-rabbit Alexa Fluor 488 (Cat# A32790, RRID:AB_2762833), for MUC5AC donkey anti-mouse Alexa Fluor 647 (Cat# A-31571, AB_162542) and for CRTAC1 donkey anti-rabbit Alexa Fluor 555 (Cat# A-

31572, RRID:AB_162543, Thermo Fisher Scientific, Waltham, MA) secondary antibodies were incubated in blocking solution for two hours at room temperature in the dark. Nuclei were counterstained with Hoechst 34580 (Cat# 565877, RRID:AB_2869723, BD Biosciences, San Jose, CA). Prolong gold mounting medium (Cat# P36934, RRID:SCR_015961, Thermo Fisher Scientific, Waltham, MA) was used for mounting. Images were acquired with an upright LSM 900 Axio Examiner 2.1 Airyscan 2 imaging system (Carl Zeiss, Oberkochen, Germany) using Zen Blue software (RRID:SCR_013672, Carl Zeiss, Oberkochen, Germany) and processed using Imaris software, version 9 (Oxford Instruments, Abingdon, U.K.)

Supplementary Figure legends.

**Supplementary Figure S1. Absolute quantification of mucin-related proteins. A:** Mass spectrometry (MS) protocol for absolute and relative quantification of proteins in bronchoalveolar lavage (BAL) fluid. **B:** Absolute quantification of mucin-associated proteins by mass spectrometry in Stable and early CLAD (eCLAD) patients at 1 month (1M) and 12 months (12M) after lung transplantation. Mucin related protein levels are juxtaposed to a reference cohort of non-smokers (NS), asymptomatic smokers (AS) and patients with chronic obstructive pulmonary disease (COPD) from a previous study (11). For comparisons between paired measurements Wilcoxon matched pairs signed rank test was used; for comparisons between two independent measurements Mann-Whitney test was used; for three independent group comparisons, Kruskal-Wallis test with Dunn's multiple comparisons test was used.. *$p < 0.05$, **$p < 0.01$, and ***$p < 0.001$ for significance.

**Supplementary Figure S2. Proteomic profile analyzed in relation to single or double lung transplant. A:** Principal component analysis of Stable and early CLAD (eCLAD) groups at 1 month (1M) and 12 months (12M), indicating single and double lung transplants (LTx). **B:** Significantly altered 23 overlapping proteins between Stable and eCLAD groups shown by heatmap, indicating single and double LTx.

**Supplementary Figure S3. Network analysis. A:** Core pathways resulting from Ingenuity Pathway Analysis (IPA). Gene ontology pathways resulting from differential proteins in bronchoalveolar lavage fluid (BALF) from Stable group at 1 month (1M) versus 12 months (12M) **(B)**, and Stable at 12M versus early CLAD group (eCLAD) at 12M **(C)**.

**Supplementary Figure S4. Protein levels of selected mediators.** Significantly changed protein levels by mass spectrometry analysis of bronchoalveolar lavage fluid (BALF) from Stable and early CLAD (eCLAD) groups at 1 month (1M) and 12 months (12M) after lung transplantation. Protein levels are juxtaposed to a reference cohort of non-smokers (NS), asymptomatic smokers (AS) and patients with chronic obstructive pulmonary disease (COPD) from a previous study (12). For comparisons between paired measurements Wilcoxon matched pairs signed rank test was used; for comparisons between two independent measurements Mann-Whitney test was used; for three independent group comparisons, Kruskal-Wallis test with Dunn's multiple comparisons test was used.*$p < 0.05$, **$p < 0.01$, ***$p < 0.001$ and ****$p < 0.0001$ for significance.

Supplementary table Legends

**Supplementary Table S1.** Isotopically labelled peptides used for absolute quantification of selected proteins.

**Supplementary Table S2.** Absolute quantification of selected proteins, protein amounts are presented as average of 1-8 peptides fmol/µl of sample.

**Supplementary Table S3.** All quantified protein groups, Log2 value for different comparisons together with p-value, Log10 LFQ values for individual samples.
